# Acquisition and extinction of drug-context memories are linked to distinct epigenetic and transcriptional mechanisms in the mouse dentate gyrus

**DOI:** 10.64898/2026.03.02.708528

**Authors:** Madelyn R. Baker, Rose Sciortino, Caitlin Zarley, Diego Scala-Chavez, Paul Bergin, Anjali Rajadhyaksha, Miklos Toth

**Affiliations:** Department of Pharmacology, Weill Cornell Medicine, New York, NY 10065; Neuroscience Program, Weill Cornell Graduate School of Medical Sciences, New York, NY 10065; Division of Pediatric Neurology, Department of Pediatrics, Weill Cornell Medicine, New York, NY 10065; Feil Family Brain and Mind and Research Institute, Weill Cornell Medicine, New York, NY 10065; Center for Substance Abuse Research, Department of Neural Sciences, Lewis Katz School of Medicine, Temple University, Philadelphia, PA 19140

**Author notes:** Correspondence should be addressed to MT, Weill Cornell Medicine, 1300 York Ave, New York, NY 10065 or AR Lewis Katz School of Medicine of Temple University, 3500 N Broad Street, MERB 852, Philadelphia, PA 19140.

## Abstract

Acquisition and extinction of drug-context associations both involve learning, yet whether extinction erases the original drug memory remains unresolved. As learning is associated with epigenetically mediated transcriptional plasticity, we asked whether acquisition-induced DNA methylation and gene expression changes are reversed by extinction, or whether extinction induces its own distinct methylation and transcriptional changes. Here, we show that both acquisition and extinction of cocaine conditioned place preference (CPP) preferentially hypomethylated cis-regulatory elements and upregulated transcription, but at largely non-overlapping genomic regions and genes in the dorsal dentate gyrus, a key region in contextual learning. In both learning paradigms, the number of differentially expressed genes was an order of magnitude smaller than those differentially methylated, highlighting the robustness of the transcriptional network to epigenetic modifications, and implicating a non-linear relationship between regulatory elements and transcription characteristic for gene regulatory networks (GRNs). Notably, animals that failed to extinguish cocaine CPP displayed attenuated DNA methylation changes and minimal transcriptional response, consistent with the stochastic output of GRNs to produce alternative outcomes across individuals. Acquisition-upregulated genes were enriched in neuronal cilium functions, consistent with the known role of primary cilia in hippocampal learning and the persistence of drug-context memories through stable axo-ciliary signaling. In contrast, extinction-upregulated genes were overrepresented in mitochondrial energy homeostasis functions, suggesting their role in meeting rapid energy demands during learning. Overall, acquisition and extinction engage fundamentally distinct molecular mechanisms, providing a potential mechanistic explanation for why drug-context memories are suppressed but not erased by extinction.

## Introduction

A central question regarding maladaptive memories and their extinction is whether extinction represents modification of the same memory or the formation of a new inhibitory memory. Among maladaptive memories, drug-context associative memories are particularly persistent and prone to relapse, highlighting the clinical importance of understanding the molecular and circuit mechanisms underlying these processes^1^.

Drug-context associative memories are formed between the rewarding effect of drugs, such as cocaine, and the spatial context in which it is experienced. For example, repeated cocaine administration in a specific environment in the conditional place preference (CPP) model leads to a preference for the now drug-associated context^2–4^. This preference can be reduced through extinction, achieved by repeated exposure to the conditioned context in the absence of the drug^3, 5^. Although extinction involves contextual learning, it does not erase the original cocaine-context memory; rather, it establishes new inhibitory learning that suppresses the original memory^6–9^. This distinction is also reflected in their durability: drug-context memories are robust and long-lasting, whereas extinction memories are more fragile and susceptible to spontaneous recovery or reinstatement by drug-associated cues or the passage of time^10, 11^. These dynamics are thought to contribute directly to relapse after abstinence^12–14^.

Aligned with its well-established role in encoding contexts and experiences, the hippocampus is a critical brain region for cocaine-context associative learning^15, 16^. Within the trisynaptic circuit of the dorsal HPC, dentate granule cells in the dentate gyrus (DG) act as a gateway for information flow from the entorhinal cortex^18, 19^. Dentate granule cells encode the cocaine-paired context and modulate, via the HPC circuit, the activity of nucleus accumbens medium spiny neurons^17^. The DG is essential for both acquisition and extinction of cocaine CPP^9, 20^. It is particularly important for context discrimination^21, 22^, and prior work has shown that activation of calcium and glutamatergic pathways in DG neurons is recruited for extinction of cocaine CPP^23, 24^. Together, these findings indicate that the dorsal DG contributes to both the formation and suppression of drug-context memories.

At the molecular level, DNA methylation and other epigenetic modifications regulate synaptic plasticity, a key substrate for learning and memory. While prior studies have examined DNA methylation in hippocampal learning, particularly contextual fear conditioning^25–29^, less is known about its role in drug-context associative learning and extinction. We asked whether acquisition of cocaine CPP is associated with specific regional DNA methylation signatures in the dorsal DG that are reversed by extinction, or whether acquisition and extinction induce distinct methylation changes.

We found that both acquisition and extinction are associated with widespread DNA methylation changes across thousands of small genomic regions, yet these changes are largely non-overlapping and exhibit distinct epigenetic plasticity. Consistent with this pattern, acquisition and extinction also induce gene expression differences in completely distinct sets of genes. Together, these findings suggest that drug-context association and extinction learning in the dorsal DG are epigenetically and transcriptionally separate processes, providing a mechanistic explanation for why extinction does not erase the original memory but instead establishes a new inhibitory memory trace.

## Materials and Methods

### Animals

All procedures followed Weill Cornell Medicine IACUC guidelines (protocol 2012-0022). Adult (8 weeks old) male C57BL/6 mice (Jackson Laboratory) were group housed (4–5/cage) in a climate-controlled facility on a 12 h light/dark cycle (6 AM–6 PM) with ad libitum food and water.

### Drugs

Cocaine Hydrochloride (Sigma Aldrich) was dissolved in 0.9% saline (1 mg/mL) and delivered intraperitoneally at 10 mg/kg (0.01 ml/g of body weight).

### Cocaine conditioned place preference (CPP)

CPP was performed as previously published^23, 24^.

### DNA Methylation Profiling by eRRBS

Genomic DNA was isolated from microdissected dorsal dentate granule cell layers and processed for enhanced reduced representation bisulfite sequencing (eRRBS) as previously reported^30, 31^.

### Bulk RNA-sequencing

Dorsal dentate gyrus tissue was microdissected from frozen brain sections, and RNA was extracted and sequenced using standard poly(A)-selected bulk RNA-seq. Reads were aligned to the mouse genome and quantified using established pipelines. Differential expression analysis was performed with DESeq2 after low-count filtering and outlier removal; significance was defined by adjusted *p* < 0.1 and |log FC| > log (1.3).

### Single nuclei RNA-sequencing

Following CPP extinction, dorsal hippocampi were isolated and nuclei were prepared using a commercial neuronal nuclei isolation kit with minor modifications. PROX1+ nuclei were labeled and purified by fluorescence-activated nuclei sorting. Sorted nuclei were processed for single nucleus (sn) RNA-seq using the 10x Genomics Chromium platform as previously published^30^.

### Statistics

Statistical analyses for behaviors were done in GraphPad Prism. Behavioral data was analyzed with a two-way ANOVA and Šidàk’s multiple comparison test. MethylKit was used to perform differential methylation and statistical analyses for differential methylation ^32^. Differentially expressed genes were determined using Benjamini-Hochberg corrected p-value. Significance of overlaps was determined using hypergeometric testing (*phyper* function in R).

## Results

### Uniform acquisition but individually variable extinction of cocaine conditioned place preference in mice

Mice were trained and tested for cocaine CPP acquisition as outlined in **Figure 1A**. Male mice exhibited robust acquisition of cocaine CPP when tested 24 hours later (day 5) in the absence of the drug, with significantly higher preference score for the cocaine-paired context compared to the baseline test and saline controls (**Figure 1B**). Virtually all animals developed preference (>90%).

**Figure 1.**
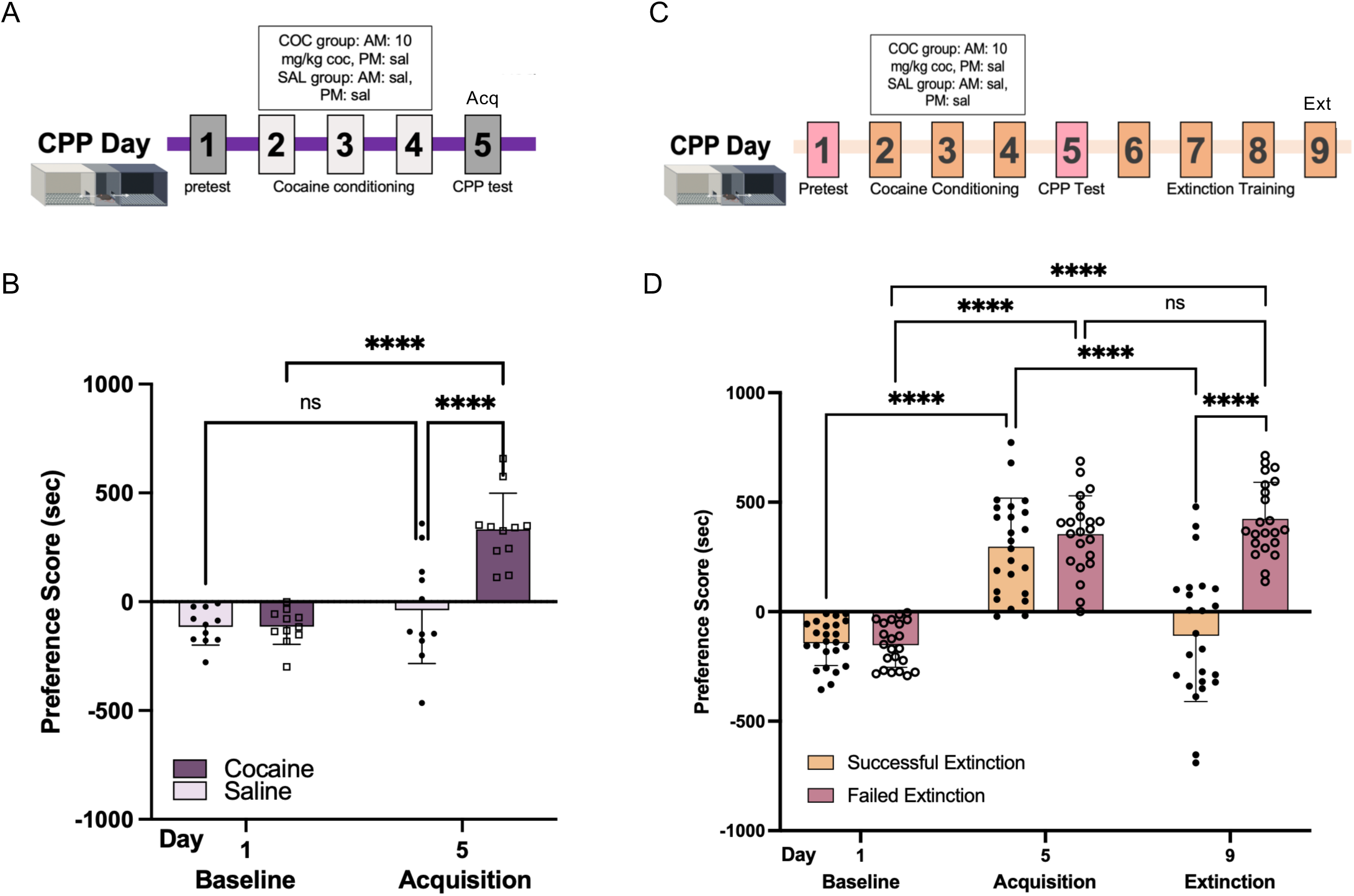
Acquisition of cocaine CPP and its extinction. **A**. Experimental timeline of the CPP protocol with 3 days acquisition (days 2-4) followed by testing on day 5. Cocaine was administered at AM, while saline at PM. **B**. Increased preference of cocaine injected mice to the drug-paired compartment of the three-chamber box on day 5. Two-way ANOVA: main effect of group (F_1,40_ = 15.03, p = 0.0004), main effect of day (F_1, 40_ = 29.84, p < 0.0001),. Šidàk’s multiple comparison test, **** p ≤ 0.0001. **C**. Experimental timeline of cocaine CPP (days 2-5) with extinction (days 5-8) and extinction testing (day 9). **D**. 52% of the animals exhibited a significant reduction (more than 70%; in preference score on day 9 (Successful Extinction), while 48% had a preference score similar to their CPP score (less than 70% reduction; and were considered resistant to extinction (Failed Extinction). Successful and Failed Extinction groups had comparable CPP preference scores in day 5. Repeated measures two-way ANOVA: main effect of group (F_1, 44_ = 21.93, p < 0.0001), main effect of day (F_1.55, 77.22_ = 100.70, p < 0.0001), interaction (group x day) (F_2, 88_ = 38.21, p < 0.0001). Šidàk’s multiple comparison test, **** *P* ≤ 0.0001.

In a second cohort, male mice underwent cocaine CPP acquisition followed by four consecutive days of drug-free extinction training (**Figure 1C**), as we have previously reported^24^. Unlike the consistency in acquisition in the group, extinction revealed substantial inter-individual variability. To operationalize extinction performance, we quantified each animal’s change in preference relative to its day 5 acquisition score. Mice that reduced their preference to <30% of their day 5 value (i.e., >70% reduction) by day 9, representing ∼52% of the cohort, were classified as exhibiting successful extinction (S-Ext, **Figure 1D, Supplementary Figure 1**). The remaining ∼48% failed to meet this criterion, maintaining ≥30% of their day 5 preference and were classified as failed extinction (F-Ext). Longer extinction, up to 7 days, was still insufficient to significantly reduce the preference of F-Ext animals indicating a true insensitivity to extinction. Females had a similar fraction of F-Ext individuals under the same conditions indicating that variability in extinction is not sex specific (**Supplementary Figure 2**). Overall, virtually all animals learned to associate the drug with a specific context but not all of them were able to learn to suppress this memory when the context was no longer associated with the drug.

### Acquisition and extinction preferentially hypomethylate CG sites in dentate granule cells

Both acquisition and extinction of cocaine CPP involve contextual learning^9, 20, 33, 34^ and hippocampal learning is associated with alterations in DNA methylation^25–29^. To determine how DNA methylation dynamics differ across phases of cocaine CPP learning, we performed genome-wide DNA methylation profiling of dorsal dentate granule cells at genomic regions enriched for CGs using enhanced reduced representation bisulfite sequencing (eRRBS ^35^) with paired end reads in five groups: cocaine acquisition (Coc-Acq), saline controls for acquisition (Sal-Acq), successful extinction (S-Ext), failed extinction (F-Ext), and saline controls for extinction (Sal-Ext).

Following the acquisition test (i.e., 24 h after the last cocaine administration on day 5, **Figure 1A**), the granule cell layer of the dorsal dentate gyrus, containing the cell bodies and nuclei of dentate granule cells, was microdissected from cryostat-cut coronal sections for methylation profiling. Using three to four biological replicates (i.e., individual mice), we profiled the methylation of 3,281,215 CG sites across all samples at 10X coverage and identified 68,238 CGs with statistically significant (q-value ≤ 0.01) methylation changes between the Coc-Acq and Sal-Acq groups (**Figure 2A**). To reduce the rate of false discovery^36^, we applied a +/- 10% methylation-change threshold, yielding 43,448 differentially methylated sites (Acq-DMSs with ≥10% change and q-value of ≤0.01)(**Figure 2A**). Methylation levels between individual mice within the Coc- Acq and Sal-Acq groups were highly correlated (r2 = 0.93-0.96) indicating that the difference between the two groups was not due to within-group variability in methylation (**Supplementary Figure 3**).

**Figure 2.**
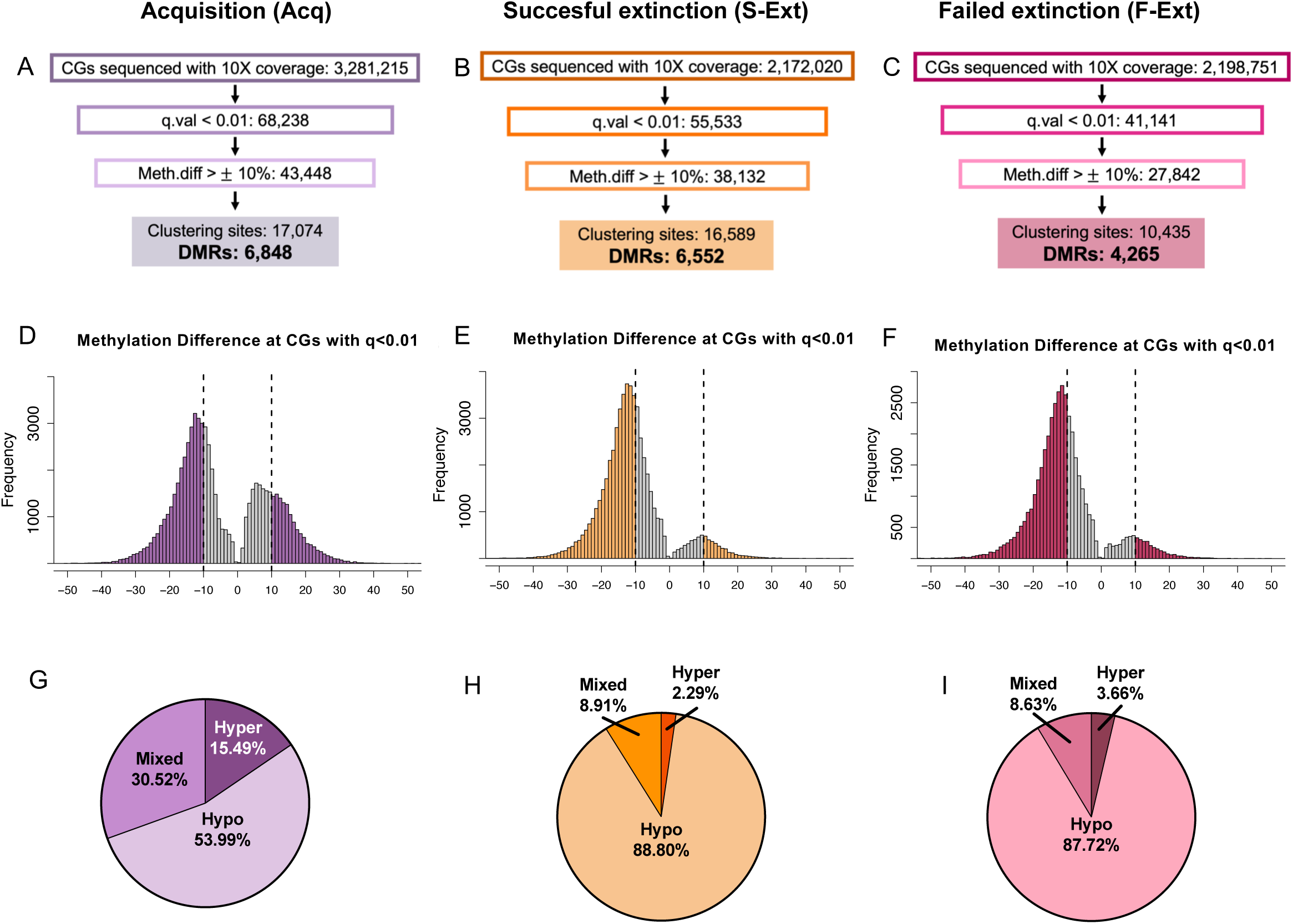
Workflow of methylation profiling and identification of differentially methylated CG sites (>10%, q<0.01) and differentially methylated regions (DMRs) in dDGCs following acquisition, and successful and failed extinction. **A**. Acq vs Saline. **B**. Successful extinction vs. saline extinction. **C**. Failed extinction vs. saline extinction. **D**. Distribution of methylation differences between cocaine Acq and saline (cocaine Acq minus saline) of CGs with 10X coverage and q-value ≤ 0.01. **E, F**. Same as (D) but between S-Ext or F-Ext and saline extinction, respectively. **G, H, I**. Fractions of Acq-DMRs, S-Ext-DMRs, and F-Ext DMRs that contain all hypermethylated sites, all hypomethylated sites, or both hyper-and hypomethylated sites (mixed).

Majority of DMSs (69.05%) were hypomethylated after acquisition (relative to Sal controls), while the rest were hypermethylated, demonstrating a hypomethylation-biased signature of acquisition (**Figure 2D**). Change in methylation at individual DMSs ranged from the 10% threshold to around 30% (**Figure 2D**), indicating that only a fraction of DMS epialleles switched from the methylated to the unmethylated state (hypomethylation) or vice versa (hypermethylation) during acquisition. Owing to the sparse sequencing coverage of eRRBS, reads represent individual cells and thus the 10-30% shift in bulk methylation level at a given epiallele can be extrapolated to a similar proportion of cells that underwent methylation switching.

Parallel methylation profiling with four replicates from individual S-Ext, F-Ext, and Sal-Ext mice following the final extinction session (day 9; **Figure 1C**) identified 38,132 S-Ext DMSs (S-Ext vs. Sal-Ext) and 27,842 F-Ext-DMSs (F-Ext vs. Sal-Ext; ≥10%, q<0.01) that were in the same range than Acq-DMSs (**Figure 2B, C**). However, the hypomethylation bias was markedly more pronounced in successful extinction (93.75%) and failed extinction (92.02%) than following acquisition, indicating an almost exclusive loss of methylation at S-Ext and F-Ext-DMSs, independently of whether extinction was successful or failed (**Figure 2E, F**). As with acquisition, change in bulk methylation level at individual DMSs after extinction ranged from the 10% threshold to around 30% (**Figure 2E, F**), indicating that extinction, whether successful or failed, involves epiallelic switching (mostly from methylated to unmethylated) in a similar fraction of cells.

Next, we clustered ≥2 DMSs within 1kb from each other into differentially methylated regions (DMRs) to assess local methylation changes in relation to gene and chromatin features. Acquisition and successful extinction were associated with comparable numbers of DMRs (6,849 and 6,553, respectively), whereas failed extinction showed somewhat fewer (4,266) (**Figure 2A-C, Supplementary Table 1A-C**). These groups also displayed similar average DMR lengths (200.4, 203.9, and 199.78 bp) and average DMS content per DMR (2.49, 2.53, and 2.45). Further, consistent with the more pronounced hypomethylation bias following extinction than acquisition, close to 90% of S-Ext and F-Ext-DMRs, while only 54% of Acq-DMRs were composed entirely of hypomethylated DMSs, with smaller proportions comprised of fully hypermethylated and mixed hypo-/hypermethylated regions (**Figure 2G-I**).

### Acquisition and extinction target genomic regions with distinct methylation signatures

DMSs within Acq-DMRs in saline controls exhibited a prominent peak between 90-100% and a smaller peak between 0-25% methylation, indicating that acquisition-sensitive sites are either methylated or unmethylated across most dentate granule cells (**Figure 3A**, light purple). Since only a relatively small fraction of Acq-DMSs displayed intermediate methylation (between 30 and 80%) in saline controls, the overall distribution resembled the canonical bimodal CG methylation landscape observed outside Acq-DMRs (**Figure 3B**) and in the mouse genome in general^37, 38^. CGs within Acq-DMRs that did not reach the 10% cutoff for qualifying as DMSs (non-DMS CGs, 7.07 per DMR on average) showed a similar, largely bimodal methylation distribution (**Supplementary Figure 5A**). As acquisition-related de novo methylation or active demethylation occurred in only ∼10-30% of cells (**Figure 2D**), the resulting bulk methylation within Acq-DMRs shifted toward intermediate values (**Supplementary Figure 4A, B**), with this pattern most evident when changes were plotted in 10% bins (**Figure 3C**).

**Figure 3.**
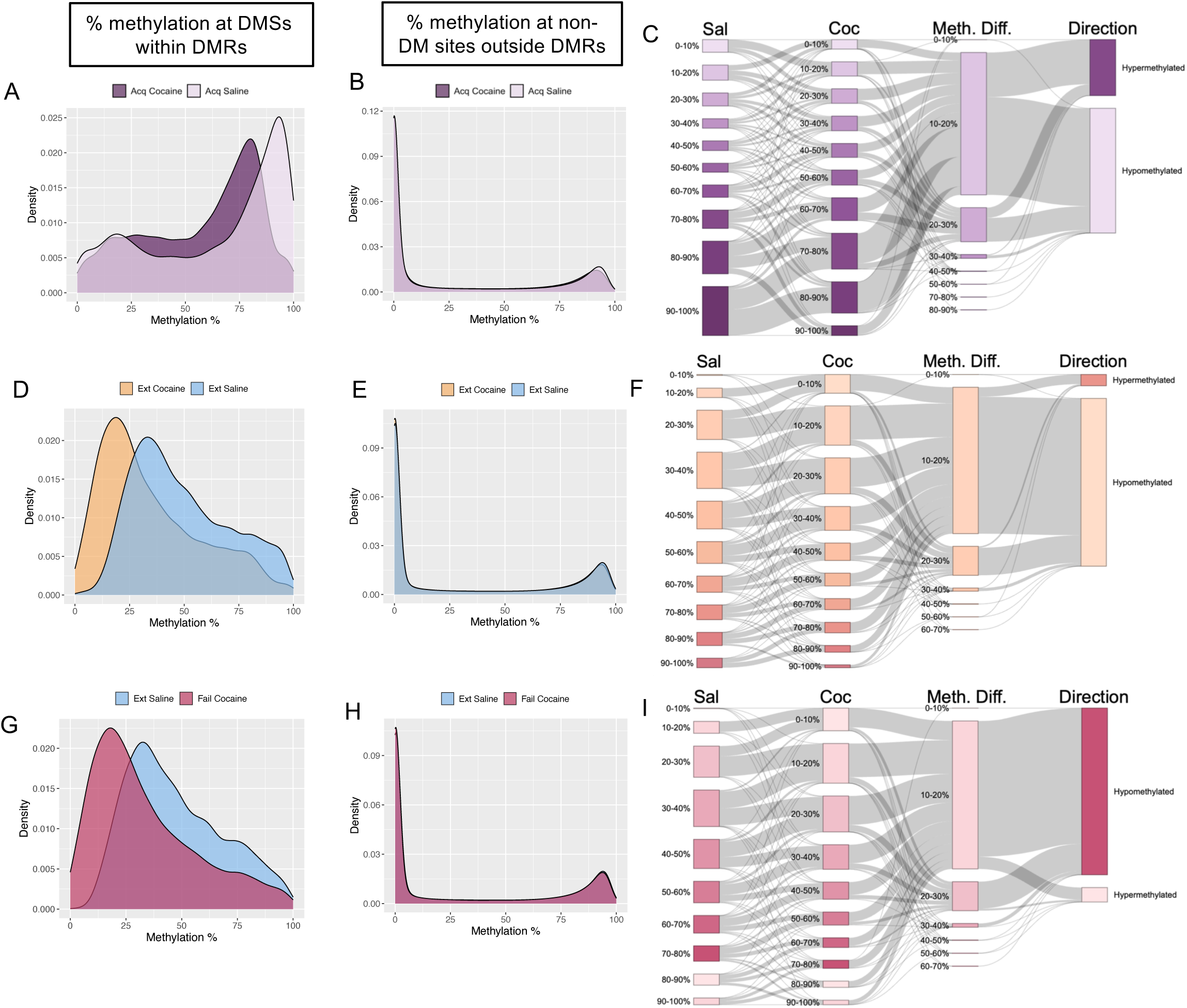
Acquisition and extinction preferentially hypomethylate CG sites in dentate granule cells. **A, D, G**. Methylation percent distribution of DMSs within Acq-DMRs for saline (light purple) and cocaine (dark purple) groups (A), DMSs within S-Ext-DMRs for S-Ext (orange) and saline (blue) groups (D), and DMSs within F-Ext-DMRs for F-Ext (pink) and saline (blue) groups (G). **B, E, F**. Methylation percent distribution of CGs outside of Acq-DMRs, S-Ext DMRs, and F-Ext DMRs, respectively, demonstrating a bimodal methylation distribution. **C, F, I**. Sankey diagram for methylation switching of DMSs within Acq-DMRs, S-Ext-DMRs, and F-Ext-DMRs from baseline in saline to Acq, S-Ext, and F-Ext DGCs in 10% increments.

In contrast, extinction targeted genomic regions with a markedly different baseline methylation profile. DMSs within both S-Ext and F-Ext-DMRs exhibited prominent intermediate methylation in saline control, with a peak at the 25-50% methylation range, indicating bistable methylation (**Figure 3D, G**, blue). Since the majority of reads in bulk eRRBS correspond to individual cells, it follows that S-Ext and F-Ext DMSs were methylated in 25- 50% of control dentate granule cells while unmethylated in the rest. CGs outside Ext-DMRs (S-Ext and F-Ext) showed the expected genome-wide bimodal methylation (**Figure 3E, H**). Methylation of non-DMS CGs within Ext-DMRs were also largely bimodal in saline control (**Supplementary Figure 5B, C**; average of 3.89 and 4.20 non-DM CGs per S-Ext-DMR and F-Ext-DMR, respectively) indicating that within DMRs, bistable DMSs are interspersed with unistable CGs. Finally, consistent with the dominant hypomethylation bias of extinction (**Figure 2H, I**), methylation of both S-Ext- and F-Ext DMSs shifted toward lower levels (**Supplementary Figure 4C, E**, mustard and dark pink), with only a minority of sites exhibiting hypermethylation following extinction (**Supplementary Figure 4D, F**). These data indicate that clusters of intermediately methylated CGs are sensitive to extinction, whether successful or failed, and that demethylation of the small, methylated fraction of these DMSs produces a nearly uniform unmethylated pattern. These extinction-related methylation shifts in S-Ext- and F-Ext-DMSs were most discernible when visualized in 10% methylation bins (**Figure 3F, I**). Overall, we conclude that although both acquisition of cocaine CPP and its extinction are predominantly associated with hypomethylation, they target regions with predominantly fully methylated and intermediately methylated CGs, respectively.

Consistent with their different methylation profiles, Acq-DMRs and S-Ext-DMRs showed limited genomic overlap, sharing only 735 overlapping DMRs (∼11% of DMRs), most of which contained only a single common DMS (762 common DMSs) at the end of the DMRs (**Figure 4A**). Overall, close to 90% of DMRs were unique to either acquisition or extinction, indicating that these two processes are associated with distinct epigenetic plasticity. This aligns with behavioral evidence that extinction represents new inhibitory learning that suppresses, rather than erases, the original memory of drug-context association^7^.

**Figure 4.**
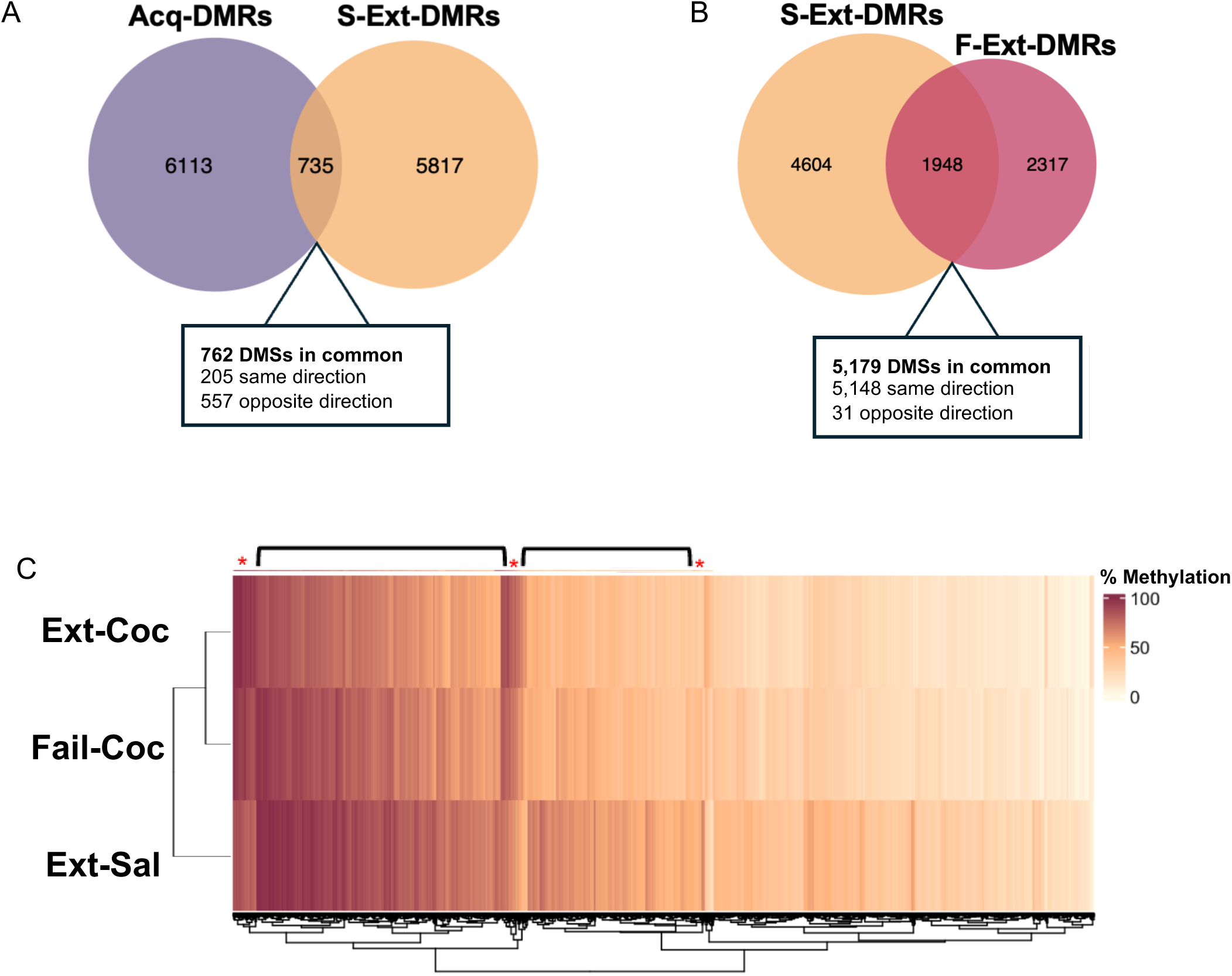
Acquisition and extinction target regions with different methylation signatures. **A.** Minimal overlap between Acq- and S-Ext-DMRs. **B**. Moderate overlap between S-Ext- and F-Ext-DMRs. **C**. Heatmap of methylation levels at S-Ext-DMR DMSs in S-Ext, F-Ext, and Ext-saline dentate granule cells.

Finally, successful and failed extinction showed more overlap in DMRs, with 30% of S-Ext-DMRs and 45% of F-Ext-DMRs shared (**Figure 4B**). Moreover, 99% of the 5,179 shared DMSs changed in the same direction, predominantly hypomethylation. Despite this, S-Ext DMRs had more total DMRs, and 70% of S-Ext-DMRs were unique to successful extinction, indicating that methylation of these DMSs might be essential for successful extinction. Closer examination of all Ext DMSs revealed that methylation changes were typically less robust in F-Ext than in S-Ext dentate granule cells, with F-Ext methylation values being between Sal-Ext and S-Ext levels and thus not significant (examples highlighted by black brackets, **Figure 4C**). The smaller subset of hypermethylated S-Ext-DMSs showed a similar pattern, with a less robust methylation shift in F-Ext dentate granule cells (examples marked by red stars in **Figure 4C**). Overall, these findings suggest that failed extinction may stem, at least in part, from an insufficient proportion of dentate granule cells switching epialleles. However, methylation changes at F-Ext-specific DMRs may also contribute to the failure extinction.

### Acquisition and Extinction DMRs are both enriched in enhancer and depleted in heterochromatin specific histone marks

The distinct methylation pattern of acquisition and extinction DMRs (i.e., mostly bimodal and intermediately methylated, respectively), suggested that they might have different chromatin states. To test this notion, genomic coordinates of DMRs were overlapped with the coordinates of publicly available chromHMM^39^ chromatin states in C57Bl/6 mouse dentate granule cells^40^. ChromHMM categorizes genomic regions into specific chromatin states based on the combination of histone modifications, including promoter, enhancer, and heterochromatin states. Acq- and Ext-DMRs had largely similar chromatin associations, but a somewhat larger fraction of S-Ext- and F-Ext-DMRs than Acq-DMRs was in a chromatin state typical for cis-regulatory elements (55% and 44%, respectively), particularly in weak promoters (H3K4me3) and active enhancers (H3K4me1 + H2K27ac) (**Figure 5A**). This difference was also reflected in the higher enrichment of Ext-DMRs than Acq DMRs in active enhancers and the selective enrichment of Ext-DMRs, but not Acq-DMRs, in weak promoters (**Figure 5B**). In contrast, a slightly larger fraction of Acq DMRs than Ext-DMRs was in facultative heterochromatin (H3K27me3), although all DMRs were relatively depleted in this widely distributed chromatin state (**Figure 5A, B**). Furthermore, Ext-DMRs but not Acq-DMRs were depleted in gene deserts. There was no apparent difference between the chromatin states of S-Ext and F-Ext DMRs (**Figure 5B**). Finally, most DMRs, both Acq and Ext, were located fully or partly in gene bodies, which include introns and exons (**Figure 5C**). Overall, these data show that both acquisition and extinction DMRs preferentially map to enhancer-associated chromatin, but at distinct genomic regions.

**Figure 5.**
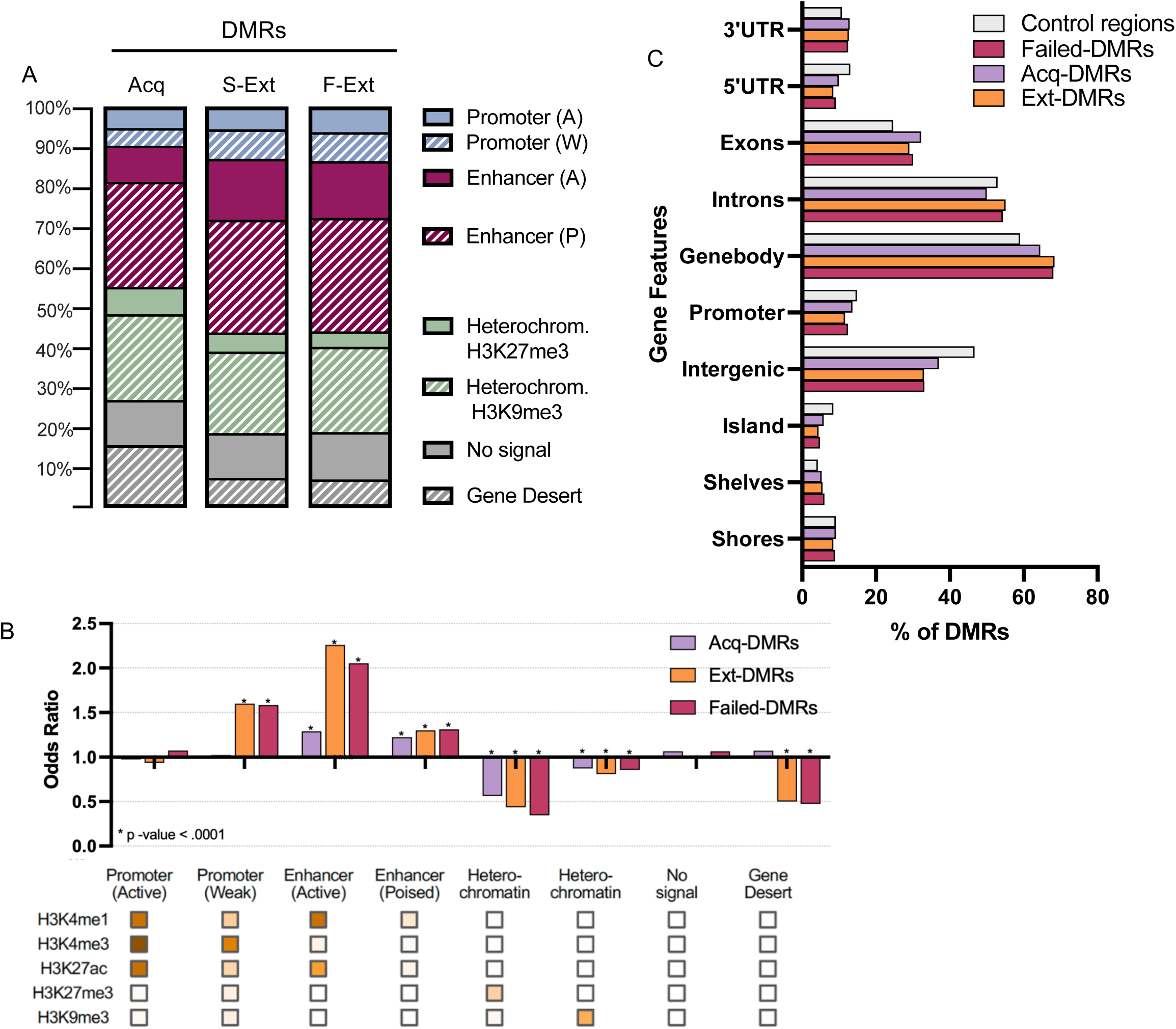
Acq- and Ext-DMRs are enriched in enhancer-specific chromatin and depleted in heterochromatin. **A.** Distribution of Acq-, S-Ext-, and F-Ext-DMRs in ChromHMM chromatin states, as determined in mouse DGCs. **B**. Enrichment/depletion of DMRs in HMM chromatin states. Enrichments were calculated using Fisher’s exact test and multiple testing correction using false discovery rate (FDR < 0.05), comparing to background block regions clustered from all tested CGs. **C**. Distribution of DMRs across gene features as compared to randomized control regions.

### Distinct transcriptomic changes following drug-context associative and contextual extinction learning

To elucidate whether acquisition- and extinction-related DNA methylation changes are correlated with gene expression, we profiled the transcriptome of dentate granule cells after acquisition and extinction of cocaine CPP. We identified 108 differentially expressed genes (DEGs) after acquisition and 280 DEGs after successful extinction (**Supplementary Tables 2A** and **5A**). The two DEG sets (Acq-DEGs and S-Ext-DEGs) showed minimal overlap (only a single gene), in line with the distinct methylation profiles associated with acquisition and extinction.

### Acquisition-differentially expressed genes are predominantly upregulated and enriched in primary cilia related biological processes

Acq-DEGs obtained by bulk RNA-sequencing of dissected dorsal dentate granule cell bodies (log2FC>1.3, adj p<0.1) were almost exclusively upregulated (106 of 108, **Figure 6A, Supplementary Table 2A**) and were significantly enriched in cilium-related GO (Gene Ontology^41^) terms (**Figure 6B**). The two “GO Biological Process” terms with the highest FDR values were “Cilium organization” and “Cilium movement,” with the remaining representing their hierarchical “parent” and “child” categories. Similarly, the top “GO Cellular Component” annotations were all cilium-associated. The primary cilium is a singular, non-motile, microtubule-based sensory antenna in neurons^42–44^ (including dentate granule cells^45–47^) that receives and transmits signals from the surrounding environment to the postsynaptic neuron^48^. Overall, 27 of the 108 upregulated DEGs (25%), were cilium related (**Figure 6C**, **Supplementary Table 2A**), including 6 cilia- and flagella-associated (*Cfap*) genes and 4 axonemal dynein heavy and light chain genes, which form the core microtubule-based structure of cilia. Using a more stringent adj. p value of <0.05, 11 of the 48 DEGs (23%) remained enriched for cilium-related processes (**Supplementary Table 2B**). Although primary cilia have been implicated in the consolidation of contextual fear in the mouse hippocampus^49^, these data show the coordinated upregulation of multiple cilium-related genes in dentate granule cells following acquisition of cocaine CPP.

**Figure 6.**
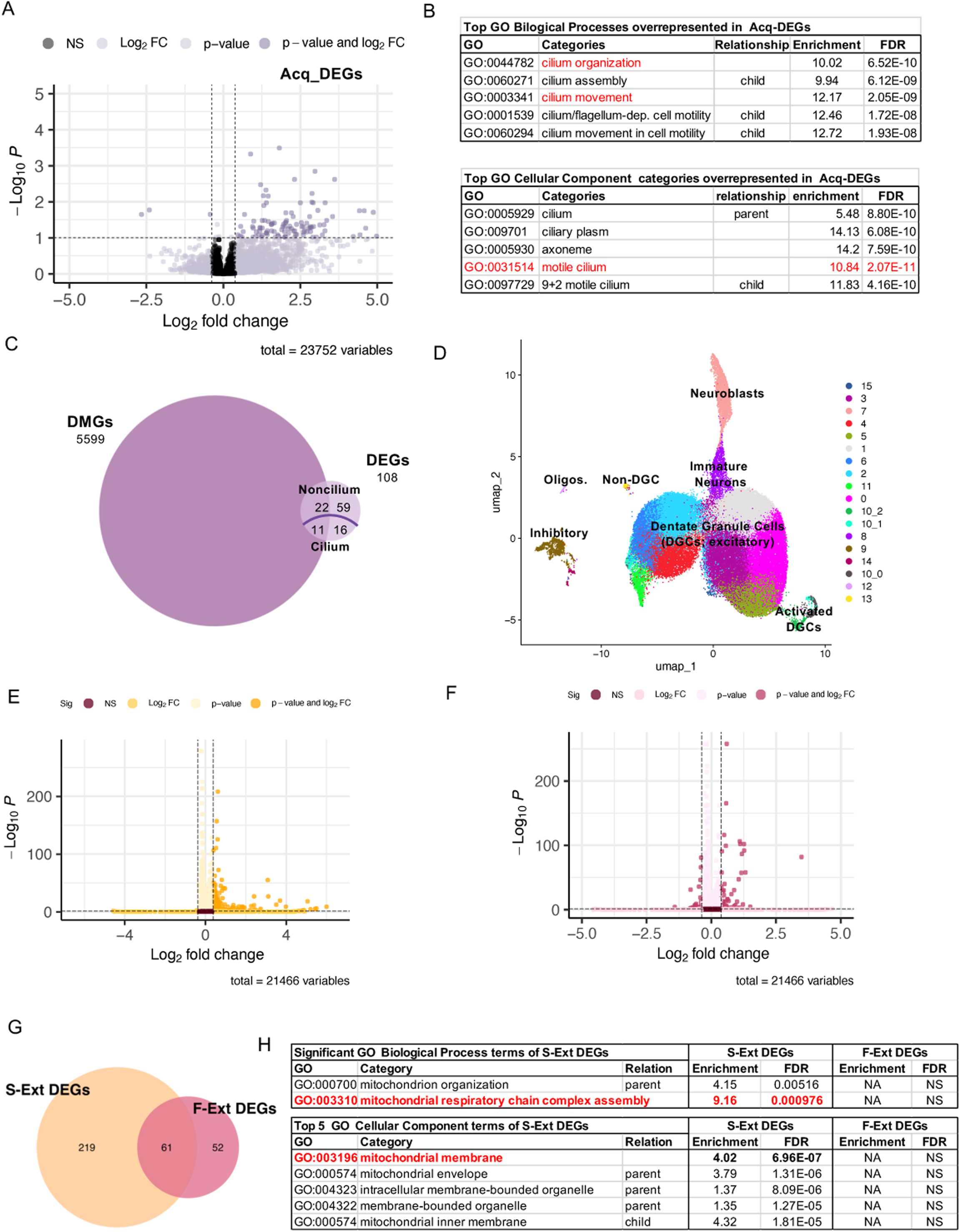
Acq-DEGs are enriched in neuronal cilium while Ext-DEGs in nuclear encoded mitochondrial respiratory complex genes. **A.** Volcano plot of differentially expressed genes by RNA-Seq from dentate granule cells following acquisition testing (Acq-DEGs). **B.** GO “Biological Processes” and “Cellular Component” overrepresentation analysis of Acq-DEGs. Red lettering indicates GO categories with the highest FDR values. **C.** Overlap between Acq-DMGs and Acq-DEGs. **D**. UMAP of Ext nuclei by snRNA-Seq. Clusters are colored and cell types indicated. **E, F.** Volcano plots of S-Ext- and F-Ext-DEGs, respectively (Wald’s test, p-adjusted < 0.1, dashed line). **G**. Overlap between S-Ext- and F-Ext-DEGs. **H**. GO “Biological Processes” and “Cellular Component” overrepresentation analysis of S-Ext-DEGs. Red lettering indicates GO categories with the highest FDR values.

To assess the relationship between differential expression and DNA methylation, we assigned DMRs to their host genes or, when located in active and poised enhancers, to their predicted target genes (i.e., differentially methylated genes or DMGs) using the “Genomic Regions Enrichment of Annotations Tool”^50^ (GREAT) (**Supplementary Table 3**). The number of Acq-DMGs (∼6,000) was considerably higher than the 108 Acq-DEGs and the overlap between the two gene sets (∼31% of all DEGs) was not statistically significant (**Figure 6C**, hypergeometric test, p=0.31). Nevertheless, the 33 Acq-DEG x DMGs (**Figure 6C, Supplementary Table 2A**) were also overrepresented in cilia-related genes (**Supplementary Figure 6, Supplementary Table 2A**). However, as with the full DEG set, cilium-specific DEGs were not differentially methylated above chance suggesting no measurable effect of acquisition-induced DNA methylation changes on their expression.

### Successful extinction is associated with the upregulation of nuclear mitochondrial respiratory chain complex genes

Because Ext-DMRs contain both methylated and unmethylated epialleles across morphologically indistinguishable dentate granule cells (**Figure 3D, G**), we expected substantial cell-to-cell variability in the transcriptional response to extinction. To capture subtle and subpopulation-specific transcriptional changes, we profiled the nuclear transcriptome of S-Ext and F-Ext dentate granule cells, along with Sal-Ext controls, using snRNA-Seq. Dentate granule cell nuclei were isolated from dorsal hippocampal nuclear preparations by fluorescence activated nuclei sorting using the granule cell marker PROX1^51^. We analyzed the transcriptomes of 26,953 S-Ext, 27,776 F-Ext, and 28,336 Sal-Ext nuclei from two replicates (5-6 mice per group in the first and 3-4 mice per group in the second replicates)^52–54^.

First, nuclei from all conditions were combined and assigned to clusters based on cell-type marker expression using Uniform Manifold Approximation and Projection (UMAP) (**Figure 6D, Supplementary Table 4, Supplementary Figure 7**). A total of 11 tightly grouped clusters represented mature dentate granule cells. Small PROX1 positive populations of interneurons and oligodendrocytes were also present, consistent with previous reports^55^, but the vast majority of transcriptionally profiled nuclei corresponded to dentate granule cells. Two small granule cell clusters (10_0 and 10_2), presumably activated during the final extinction session expressed the immediate early genes *Arc*, *Fos, and FosB* (**Supplementary Figure 7**). The individual UMAPs of S-Ext, F-Ext, and Sal-Ext nuclei were highly similar (**Supplementary Figure 8**), and the distribution of nuclei across individual clusters did not differ appreciably among groups (not shown), indicating that extinction, whether successful or failed, does not alter overall cell identity or global transcription.

To parallel the analysis of Acq-DEGs, we performed differential expression (S-Ext vs. Sal-Ext and F-Ext vs. Sal-Ext) with Seurat on all and individual mature dentate granule clusters (**Figure 6D**). Analyses on individual clusters yielded comparable but lower powered results. Analysis across all mature granule cell clusters identified 280 S-Ext DEGs and 113 F-Ext DEGs, indicating that successful extinction is associated with 2.4 times more DEGs than failed extinction (**Supplementary Table 5A, B**). As **Figure 6E** and **F** show, most Ext-DEGs were upregulated, particularly S-Ext DEGs (279 out of 280). S-Ext and F-Ext DEGs shared 61 genes that represented over half of F-Ext DEGs (**Figure 6G, Supplementary Table 5C**). Overall, successful extinction was associated with ∼4 times more unique genes (219 DEGs) than failed extinction (52 DEGs). GO analysis revealed that the full set of all S-Ext DEGs was significantly enriched only in the GO Biological Process terms “Mitochondrial respiratory chain complex assembly” and its parent “Mitochondrial organization,” whereas the 113 F-Ext -DEGs showed no significant GO term enrichment (**Figure 6H**). GO “Cellular Component” analysis similarly demonstrated enrichment of S-Ext-DEGs in mitochondrial membrane-associated annotations (**Figure 6H**). The 219 S-Ext unique DEGs displayed the same enrichment (not shown), indicating that overrepresentation of nuclear mitochondrial genes was not driven by the 61 DEGs shared by S-Ext and F-Ext. Among S-Ext DEGs, 28 genes (10% and 14% of all and unique S-Ext-DEGs, respectively) were annotated as mitochondrial and were all nuclear-encoded and included *ndufaf3, ndufs8, ndufb10, and ndufb11*, four nuclear mitochondrial NADH dehydrogenase subunit genes, several mitochondrial import receptor genes (Tomm 5, 6, and 22), import translocase subunit genes (Trimm8b, 10b, and 13), and *coa3* and *cox14*, encoding two cytochrome c oxidase assembly factors (**Supplementary Table 6**).

Of the 280 S-Ext-DEGs, only 20 (7%) were S-Ext DMGs (**Supplementary Table 7A, B**), none of which were mitochondrial genes, a frequency significantly lower than chance (significant depletion, p=2.54×10-7). This suggests that S-Ext genes are largely resistant to extinction- induced DNA methylation changes, and their differential expression may instead be mediated by other epigenetic mechanisms or by trans-regulatory inputs from other loci. A closer inspection of all and unique S-Ext DEGs revealed that their genome spans were almost an order of magnitude shorter than the genome average (**Supplementary Figure 9A, B**), which likely contributed to their depletion in DMRs and DMGs. In contrast, the size of S-Ext DMGs exceeded the genome average (**Supplementary Figure 9C**). Short genes represent a minority of the mammalian genome but play an important role in processes requiring rapid responses^56^ such as memory formation^57–59^. Overall, successful extinction DEGs were distinct from acquisition DEGs, and the upregulation of nuclear-encoded mitochondrial respiratory chain complex genes clearly distinguishes successful from failed extinction.

## Discussion

The main finding of our study is that cocaine-context learning and its extinction both reshape the DNA methylation landscape and transcriptional output of dentate granule cells, but they do so through distinct genomic targets and molecular pathways. The molecular differences mirror behavioral evidence that extinction does not erase the original memory of the drug-context association but rather suppresses it as a new, inhibitory memory trace^6–9^ that is inherently less stable and more prone to recovery^6–9^. This molecular divergence offers a potential mechanistic explanation for why drug-context associations remain persistent, whereas extinction memories are comparatively less durable and susceptible to spontaneous recovery and reinstatement.

### Acquisition and extinction of cocaine CPP target genomic regions with different baseline DNA methylation states

Both acquisition and extinction were associated with regionally clustered methylation changes (i.e., DMRs), yet Acq-DMRs and Ext-DMRs displayed fundamentally different baseline methylation states. Acquisition primarily modified CGs that were uniformly methylated across all dentate granule cells, whereas extinction targeted bistable, intermediately methylated CGs, in some but not all cells within otherwise morphologically homogenous population of mature dentate granule cells. Of note, methylation heterogeneity at extinction DMRs was unlikely due to young immature neurons in the dentate gyrus because their population is smaller than 10%, the lowest level of differential methylation considered significant in our experiments, and the 30-40% average methylation at bistable sites. Thus, acquisition and extinction engage genomic regions with substantially different initial methylation profiles.

What accounts for the regionally distinct responsiveness of the dentate methylome to the acquisition and extinction learning processes? As Ext-DMR CGs are intermediately methylated at the population level, an atypical feature in a genome largely split between uniformly methylated or unmethylated states, we propose that epigenetic metastability at bistable regions may explain their enhanced malleability during extinction learning. We recently reported similar shifts in epiallelic proportions at intermediately methylated regions in fetal ventral dentate granule cells to a proinflammatory gestational environment^30^ and in dorsal dentate granule cells of adult mice following chronic wheel running and unpredictable stress (unpublished data), suggesting that intermediately methylated regions are particularly sensitive to stimuli from the external environment. Methylation-bistable regions could function as environmental sensors and their epiallelic switching may represent adaptive or maladaptive responses. In contrast, acquisition predominantly engaged uniformly methylated and less frequently, uniformly unmethylated regions, although it also switched the methylation state of some intermediately methylated regions. Because uniformly methylated regions are typically resistant to active demethylation and unmethylated regions are protected against de novo methylation, the acquisition-specific demethylation we observed suggests that cocaine exposure perturbs ordinarily robust maintenance mechanisms. We propose that cocaine, by increasing dopamine and/or norepinephrine levels^62^, amplifies the entorhinal glutamatergic input onto granule cells and perturbs the otherwise robust epigenetic maintenance of methylated and unmethylated states. We found a similar demethylation of uniformly methylated regions following cocaine self-administration in dorsal dentate granule cells (unpublished), supporting the idea that the presence of cocaine during acquisition and its absence during extinction fundamentally shapes DMR profiles of these two learning processes.

Switching of methylation states at Acq- and Ext-DMRs was not random across the genome as cis-regulatory regions, in particular enhancers, were overrepresented among the targets suggesting that transcription factors, cofactors, and/or associated histone modifications may dictate, at least partly, their epigenetic malleability.

### Methylation state switching by acquisition and extinction involves only a subset of dentate granule cells

Acquisition and extinction shifted DNA methylation by 10-20% implying methylation-state switching in a similar proportion of cells. Whether these changes occur in overlapping sets of dentate granule cells is unknown. Given that experiences in the dentate gyrus are encoded in small neuronal ensembles (estimated to be 7-8% by calcium imaging^63^) and acquisition and extinction unfold over multiple days, it is reasonable to hypothesize that most epiallelic switching reflects cumulative population of granule cells activated throughout training. It is not known however, if extinction continues to recruit some of the acquisition activated cells or recruits a non-overlapping set of cells. The ∼20% and up to 30% methylation shifts at DMSs could correspond to the cell population that was repeatedly recruited, while lower methylation shifts may be related to cell populations that entered or left the ensemble during the trials or may represent DMSs that were less amenable for methylation switching.

### Genome-wide hypomethylation is associated with a small number of upregulated genes in both acquisition and extinction

Both acquisition and extinction were dominated by hypomethylation, and by transcriptional upregulation, raising the possibility that demethylation and gene activation are shared features of both drug-context associative and contextual extinction learning. This association was further supported by the enrichment of DMRs in promoters and enhancers, given the inverse correlation between methylation of cis-regulatory elements and gene expression. Yet, there was a striking difference that emerged between the scale of differential methylation and differential expression. Approximately 50% of the expressed genes in granule cells harbored or were associated with Acq-DMRs and S-Ext-DMRs (i.e., DMGs), while only 1-2% of genes were differentially expressed, indicating that most methylation changes were not directly associated with detectable transcriptional changes. Moreover, Acq-DEGs were not enriched in DMRs and S-Ext-DEGs were actually depleted in DMRs, likely because of their small size relative to the average size of DMGs. This may suggest that DNA methylation changes and gene expression are not linearly coupled. However, animals with failed extinction relative to successful extinction had approximately 2/3^rd^ of the DMRs and less than half of DEGs suggesting that differential methylation and expression in dentate granule cells scales with the behavioral outcome. The widespread hypomethylation (alongside limited hypermethylation) in gene bodies is reminiscent of global hypomethylation in cancer^64, 65^ and aging^66^ and suggests the reorganization of the epigenetic landscape during acquisition and extinction. Overall, our findings are inconsistent with local epigenetic regulation of DEGs and more compatible with a nonlinear relationship between the activity of cis-regulatory elements and the transcriptional output of gene regulatory networks (GRNs)^67–69^. GRNs, due to their inherent robustness, remain stable and unperturbed by manipulations but can still respond to environmental changes via highly connected nodes whose expression is adaptable^70^. The output of GRNs is difficult to predict from individual network components because regulatory elements interact in complex, non-additive ways, often involving 3D chromatin folding, cooperation, and competition for binding sites. We speculate that the small set of Acq-DEGs represent the maladaptive transcriptional output of dentate granule cells to repeated exposure to a context in the presence of cocaine, while S-Ext-DEGs correspond to the output to repeated exposure to the previously drug paired but now familiar context without the pharmacological effect of cocaine. Stochasticity is another property of GRNs providing a population with different outputs^70^ that may explain the different DNA methylation and gene expression profiles of animals with successful and failed extinction of the drug-context associative memory. Although the topology of the putative Acq and S-Ext GRNs is not known, a system level approach may help to better understand the mechanism underlying formation of the drug associated memory and the individual variations in extinction.

### Acquisition upregulates cilium specific genes while extinction upregulates nuclear mitochondrial genes in dentate granule cells

Gene ontology analysis revealed that Acq-DEGs were enriched in cilium-related functions. Primary cilia are sensory organelles, present in essentially all neurons, and central to sensing and responding to the environment^42, 43^. Although primary cilium has been reported to be required in the mouse hippocampus for the consolidation of contextual fear memory^49^, our data is the first showing the coordinated upregulation of a number of cilium genes in dentate granule cells following acquisition of cocaine CPP. Additionally, deletion of cilia in dopaminergic (DAT expressing) neurons was shown to exhibit reduced locomotor sensitization to cocaine as well as reduced CPP^71^. We identified a total of 33 cilium related genes, including cilia-associated *Cfap* genes and axonemal dynein heavy and light chain genes encoding components of molecular motors powering retrograde intraflagellar transport^72^. Given their structure and stability, axo-ciliary synapses^48^ differ from traditional axo-dendritic synapses and contribute to the formation of remote memories^49^, and thus upregulation of neuronal cilium genes in the dentate gyrus supports the persistence of drug-context associative memories.

By contrast, an entirely different set of genes were upregulated during extinction, which is consistent with the different nature of extinction and acquisition memories and in line with behavioral data showing that extinction does not erase drug-context memory. Successful extinction upregulated genes involved mitochondrial respiration and mitochondrial envelope/inner membrane compartment structure suggesting their role in energy homeostasis. Learning and memory processes rely heavily on mitochondrial energy production, including local translation, and mitochondrial respiratory function and enhanced presynaptic mitochondrial energy production have been shown to be essential for spatial remote memory and contextual fear memory formation, respectively^57–59^. Importantly, these mitochondrial genes were not upregulated in animals that failed extinction. In fact, the small set of F-Ext-DEGs were not significantly enriched in any biological function.

In summary, acquisition-driven upregulation of cilium genes may support the formation and/or stabilization of drug-context memory, while extinction-driven upregulation of nuclear mitochondrial genes may facilitate rapid encoding of a new memory.

## Supporting information

CPP extended materials and methods

## Acknowledgements

We would like to thank the support of the Epigenomics Core and the Genomics Core of Weill Cornell Medicine. We are grateful for grant support 5R01MH117004 and 5R01NS106056 to MT, R01DA053261, R01DA054368, R01MH125006 and R01DA050454 to AMR, and NSF GRFP 2139291 to MRB.

