## Supplementary material for "Acquisition and extinction of drug-context memories are linked to distinct epigenetic and transcriptional mechanisms in the mouse dentate gyrus": CPP extended materials and methods

**Cocaine conditioned place preference**

Mice underwent 7-day acclimation and 1 h daily habituation to the behavior room. CPP was performed in a three-chamber apparatus (Med Associates) using a 5-day protocol. Day 1 assessed baseline preference. On days 2–4, morning sessions paired cocaine (10 mg/kg) or saline (controls) with one chamber; afternoon sessions paired saline with the opposite chamber. Day 5 tested acquisition. Brains were collected and flash frozen. A separate group underwent extinction (days 6–9). Brains were collected following the final extinction session.

**DNA extraction and eRRBS**

Frozen brains were sectioned coronally at 200 µM slices. The dorsal dentate granule cell layer was micro-dissected. DNA extracted using the QIAamp DNA Micro Kit (Qiagen). eRRBS library preparation, sequencing, and trimming were performed by the WCM Epigenomics Core. DNA from four mice per group was sequenced (single-end 100 bp) on an Illumina NovaSeq6000.

**Differential methylation analysis**

Reads were aligned to mm10 and methylation called using Bismark. Differential methylation was analyzed with methylKit. CpGs with ≥10× coverage and present in all samples were included. Differentially methylated sites were defined as ≥10% methylation difference with q ≤ 0.01. Differentially methylated regions (DMRs) contained ≥2 differential CpGs within 1 kb and were classified as hypo-, hyper-, or mixed.

**Enrichment of DMRs in annotated chromatin states and genomic features**

DMR enrichment across chromHMM-defined chromatin states and genomic annotations (exons, introns, UTRs, promoters ±500 bp) was computed using odds ratios. Significance was assessed by χ² test with Benjamini–Hochberg correction. Control regions were generated from non-differentially methylated CpGs clustered using DMR criteria.

**RNA extraction and Bulk RNA-sequencing**

Frozen brains were sectioned (200 µm), the dorsal dentate gyrus micro-dissected, and RNA extracted using the RNeasy Mini Kit (Qiagen). Library preparation (NEB Ultra II Directional with Poly-A selection) and paired-end 2×50 bp sequencing were performed by the WCM Genomics Core (≥40M reads/sample). Reads were trimmed (cutadapt), aligned to mm10 (STAR), and quantified using Cufflinks and HT-seq^1^.

**Bulk RNA-sequencing differential expression analysis**

DESeq2 was used for differential expression. Genes with <10 counts were removed. Outliers were identified with robust PCA. Differentially expressed genes met adjusted p < 0.1 and [log₂FC] > log₂(1.3).

**Single nucleus isolation**

Following CPP Extinction and cervical dislocation, brains were extracted and cut coronally into 1mm sections on a brain block. The dorsal hippocampus was dissected from the appropriate sections and collected into D-PBS (without Ca2+ or Mg2+) with 1X protease inhibitor on ice. Before nuclei isolation, the tissue was spun down and the buffer was removed. Nuclei were isolated using the Minute™ Single Nucleus Isolation Kit for Neuronal Tissues/Cells (Invent Biotechnologies, Inc.) with slight modifications to manufacturer protocol. After adding 200 ul of Buffer A, tissue was homogenized with a pestle using 30 twists. An additional 400 ul of Buffer A was added, and tissue was homogenized with another 10 twists and then incubated at -20^o^C for 10 minutes. Homogenate was first filtered through a 70 uM filter and then transferred to a filter column provided in the kit. The homogenate was spun at 800g for 5 minutes at 4^o^C. The pellet was resuspended with 500 ul of 1% BSA in DPBS with 0.2U/ul of RNase inhibitor and spun again at 600 g for 5 minutes at 4^o^C. Nuclei were resuspended in the same buffer type to a concentration of 1000-5000 nuclei/ul.

**Fluorescence activated cell sorting**

Nuclei were stained for sorting as previously described. Briefly, first, nuclei were incubated with FC block for 10 minutes on ice. Nuclei were then stained with anti-PROX1 Alexa Flour 647 (NBP-1-30045AF647, NovusBio, 1:5000) and incubated on ice for 30 minutes. Nuclei were washed with equal volume of the appropriate buffer and spun at 600 g for 5 minutes at 4^o^C. Nuclei resuspended to a concentration of 1000-5000 nuclei/ul were sorted on a BD Influx sorter.

**Single nucleus RNA sequencing (snRNA-seq)**

Final nuclei suspensions (2 replicates per sample group – 3 groups, each pool of nuclei from 5-6 animals) were submitted to the Weill Cornell Epigenomic Core for library preparation and sequencing. Sample libraries were prepared using the Chromium Single Cell Reagent V3 Kit from 10X Genomics according to manufacturer instructions. Libraries were sequenced on an Illumina NovaSeq6000 to a depth of approximately 25,000 target reads per nuclei (approximately 250 million reads per sample of 10,000 nuclei).

**snRNA-seq analysis**

Raw sequencing data FASTQ files were processed using CellRanger v6.0.0 pipeline and aligned to the mm10 transcriptome reference with introns included. Gene expression matrices of all samples were loaded into R and assessed for quality and analyzed using Seurat^2, 3^. Nuclei with fewer than 500 genes, greater than 4,000 genes, or greater than 0.5% mitochondrial gene reads were excluded from analysis. Genes expressed in fewer than 5 nuclei were also filtered out. Normalization was performed with SCTransform with mitochondrial genes regressed out. Following PCA, clustering (using 30 PCs and resolution 0.8) was performed with FindClusters. Data was prepped with PrepSCTFindMarkers and cluster markers were found with FindAllMarkers function (min.pct = 0.2) using the default Wilcoxon rank sum test with Bonferonni correction, and cell types were annotated using known cell markers. One cluster was further clustered using FindSubCluster and a resolution of 0.1. For differential expression testing, the three groups (successful extinction, failed extinction, saline) were compared using FindMarkers with a minimum log2FC of 1.3 and adjusted p-value < 0.05.

**Gene ontology and network analysis**

Gene ontology enrichment analysis was performed using geneontology.org^4-6^. Network analysis was done using ShinyGO^7^.
