## Supplementary material for "Acquisition and extinction of drug-context memories are linked to distinct epigenetic and transcriptional mechanisms in the mouse dentate gyrus": Supplmentary Figures

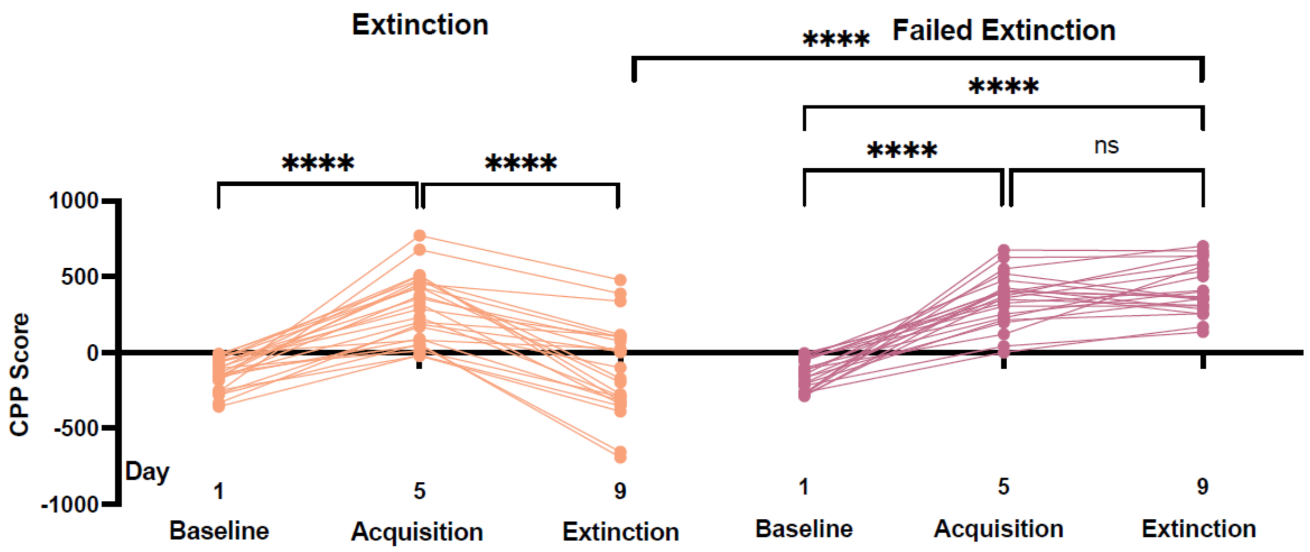


**Supplementary Figure 1**. Data presented in **Figure 1D** displayed as preference scores of individual mice through time, from baseline to acquisition and extinction. All data were analyzed together but individuals with successful and failed extinction were separated to highlight the difference in extinction between the two groups. Two way RM ANOVA, main effect of day ($F_{1.755, 77.22} = 100.70$; $p < 0.0001$), Šidàk's multiple comparison test, ****$p<0.001$; Successful extinction N=24, failed extinction N=22.

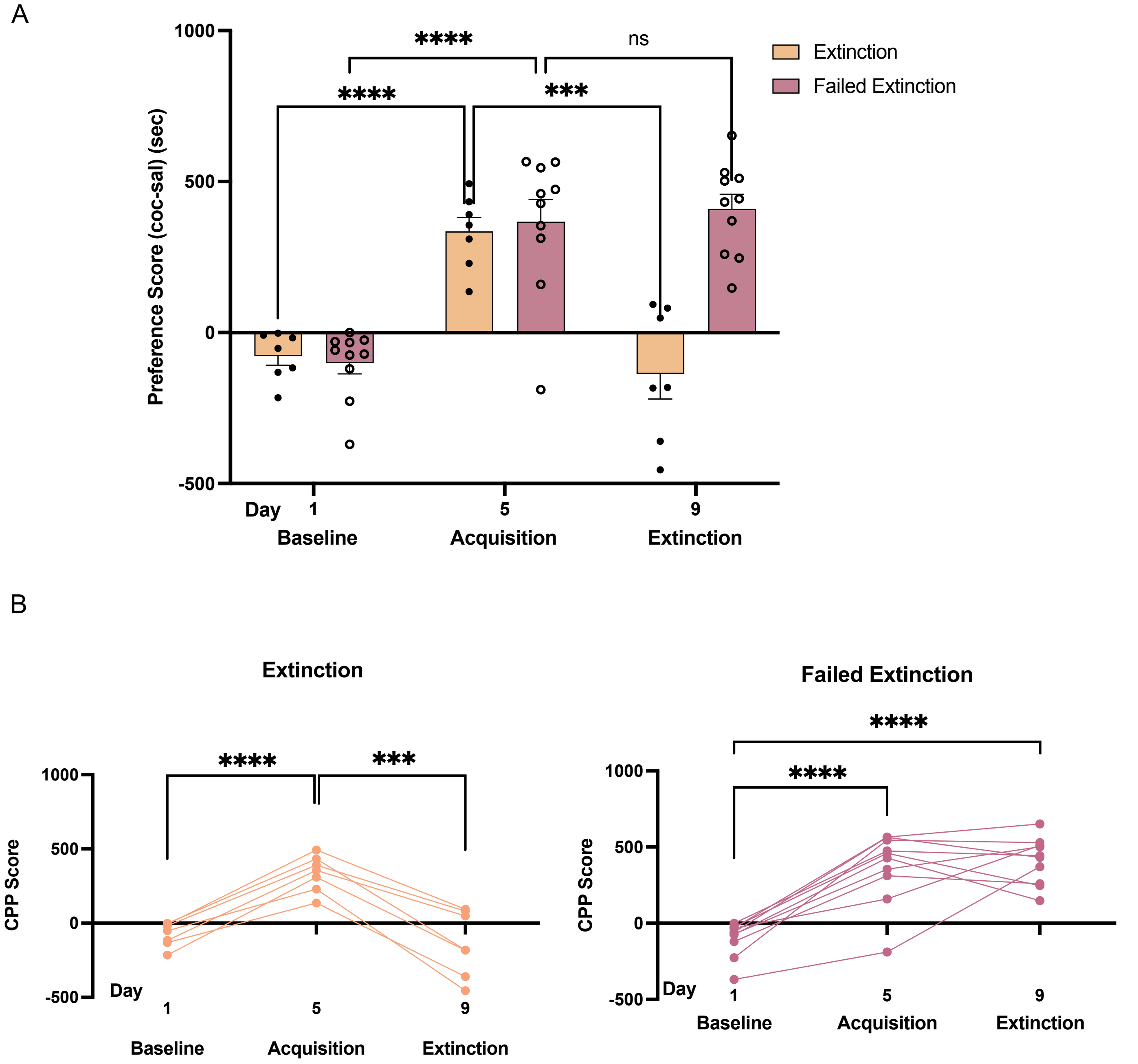


**Supplementary Figure 2**. **Extinction trials with female mice**. **A**. Individuals can be segregated to successful (over 70% reduction in day 9 vs. day 5) and failed extinction (less than 70% reduction) groups. RM two-way ANOVA: main effect of group (F1, 15 = 9.265, p =0.0082), Šidàk's multiple comparison test, ***p=0.001, ****p<0.0001. Successful extinction N=7, failed extinction N=10. **B**. As in "A", but the preference scores of individual mice are displayed through time, from baseline to acquisition and extinction. RM two-way ANOVA, main effect of day (F1.750, 26.24 = 43.38); p < 0.0001).

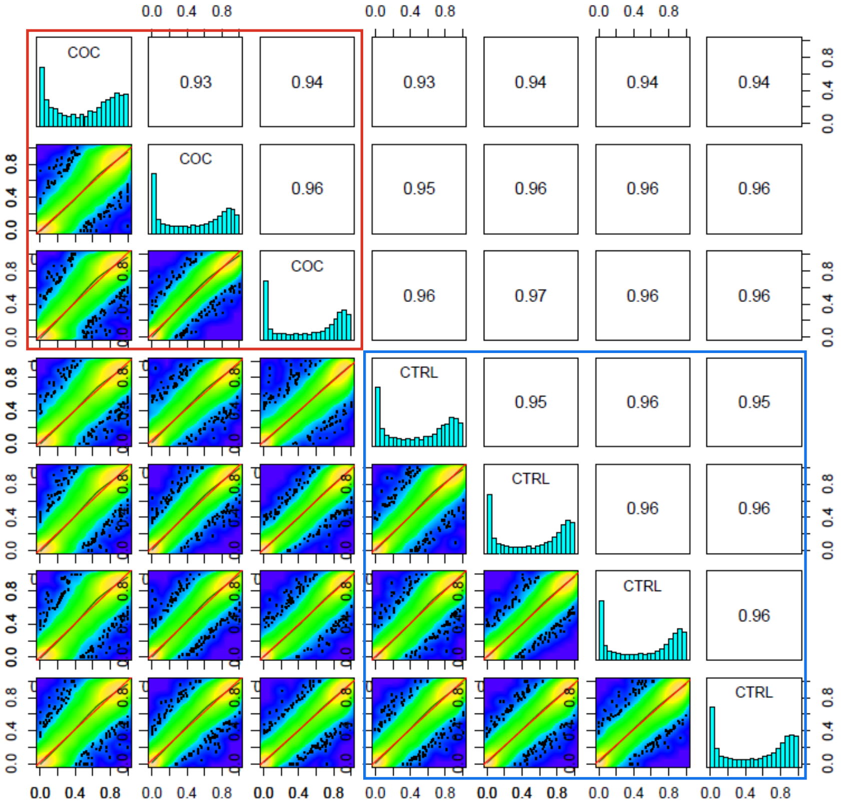


**Supplementary Figure 3**. Correlation plots of methylation at DM CGs between individual animals following cocaine (COC 1, 2, 3) and saline control (CTRL 1, 2, 3, 4) CPP. X and Y coordinates: methylated fraction. Numbers in boxes: r2. Red square includes comparisons between Coc animals, while blue square between CTRL (Sal) animals. Outside of these boxes are comparisons between individuals from the two groups.

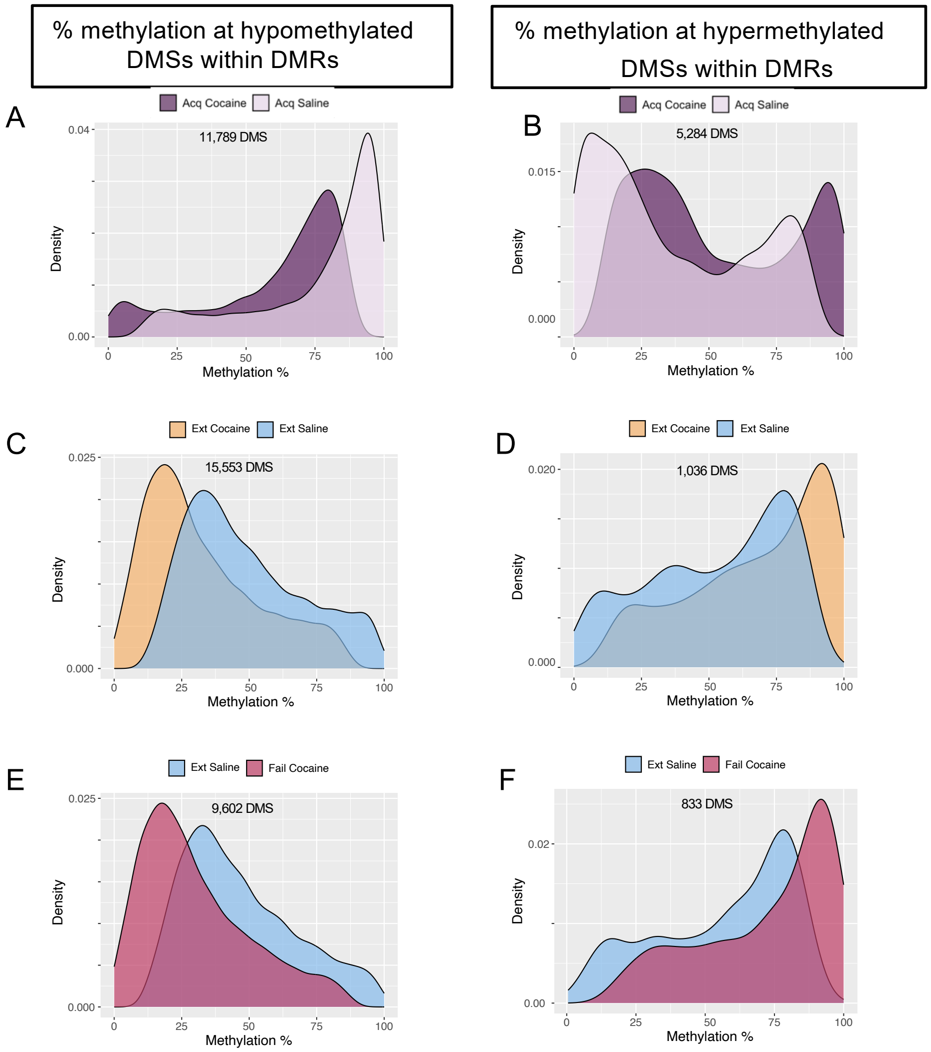


**Supplementary Figure 4**. Methylation percent distribution of clustering hypo- and hypermethylated DMSs. **A, B**. DMSs in Acq-DMRs for saline (light purple) and cocaine (dark purple) groups. **C, D**. Same as A-B but with DMSs within S-Ext-DMRs for S-Ext (mustard) and saline (blue) groups. **E, F.** Same as A-B but with DMSs within F-Ext-DMRs for F-Ext (dark pink) and saline (blue) groups.

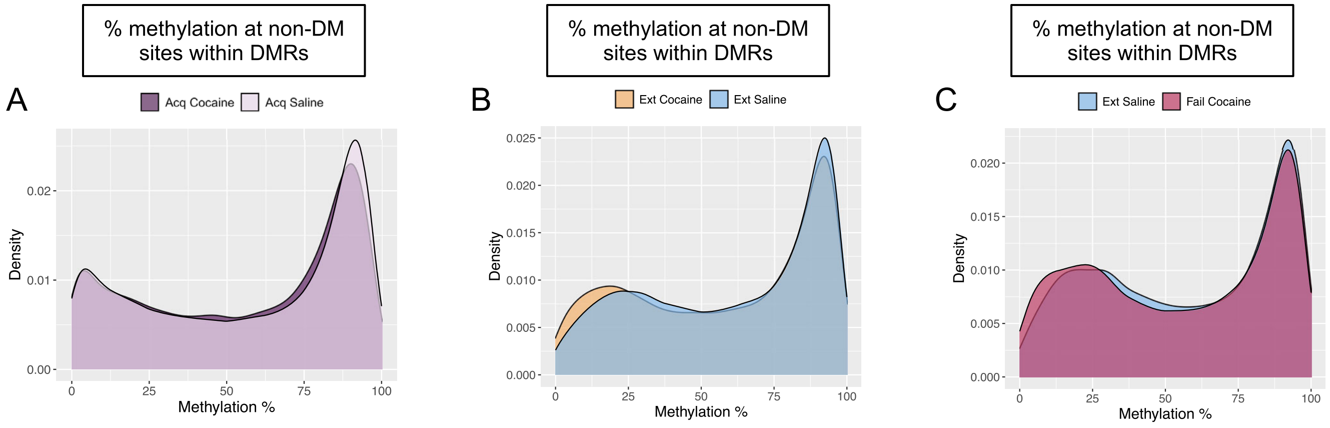


**Supplementary Figure 5**. Methylation percent distribution of non-DMS CG within DMRs that did not reach the 10% cutoff for qualifying as DMSs. **A**. Non-DMSs in Acq-DMRs for saline (light purple) and cocaine (dark purple) groups. **B**. Same as A, but within S-Ext-DMRs for S-Ext (mustard) and saline (blue) groups. **C.** Same as A but with DMSs within F-Ext-DMRs for F-Ext (dark pink) and saline (blue) groups.

| Top 5 GO Biological Processes Categories overrepresented in Acq-DMG-DEGs | | | | |
|---|---|---|---|---|
| **GO** | **Categories** | **Relationship** | **Enrichment** | **FDR** |
| GO:0044782 | cilium organization | | 11.3 | 1.70E-02 |
| GO:0060271 | cilium assembly | child | 12.52 | 1.71E-02 |
| GO0003341 | cilium movement | | 13.96 | 1.96E-02 |
| GO:0120036 | plasma membrane bounded cell projection assembly | parent | 9.79 | 2.17E-02 |
| GO:0030031 | cell projection assembly | parent | 9.52 | 2.09E-02 |

| Top 5 GO Cellular Components Categories overrepresented in Acq-DMG-DEGs | | | | |
|---|---|---|---|---|
| **GO** | **Categories** | **Relationship** | **Enrichment** | **FDR** |
| GO:0005930 | axoneme | child | 15.51 | 1.62E-02 |
| GO:009701 | ciliary plasm | | 15.44 | 1.10E-02 |
| GO:0032838 | plasma membrane bounded cell projection cytoplasm | parent | 10.46 | 4.22E-02 |
| GO:0005929 | cilium | parent | 6.04 | 2.23E-02 |
| GO:0097729 | 9+2 motile cilium | child | 11.31 | 3.65E-02 |

**Supplementary Figure 6**. Enrichment of Acq-DEG x DMGs in cilium related GO terms.

A

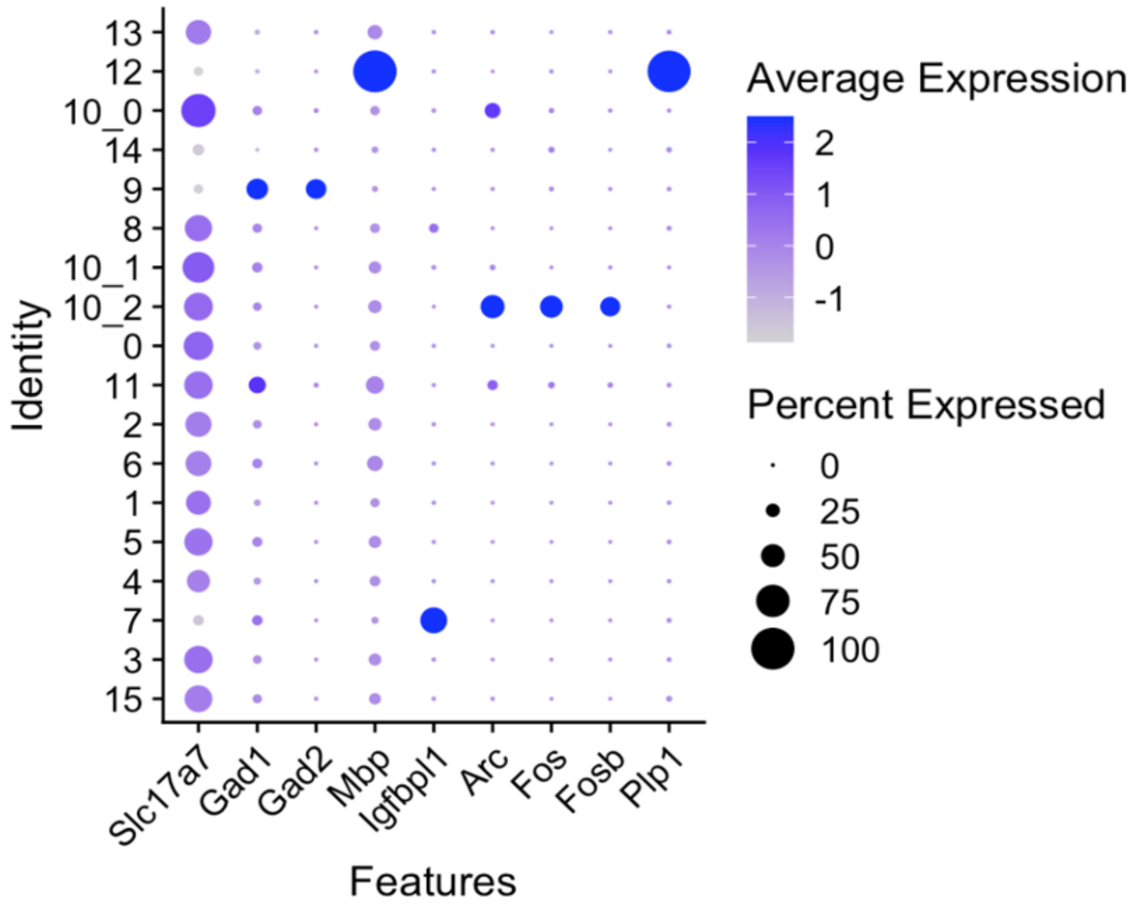


B

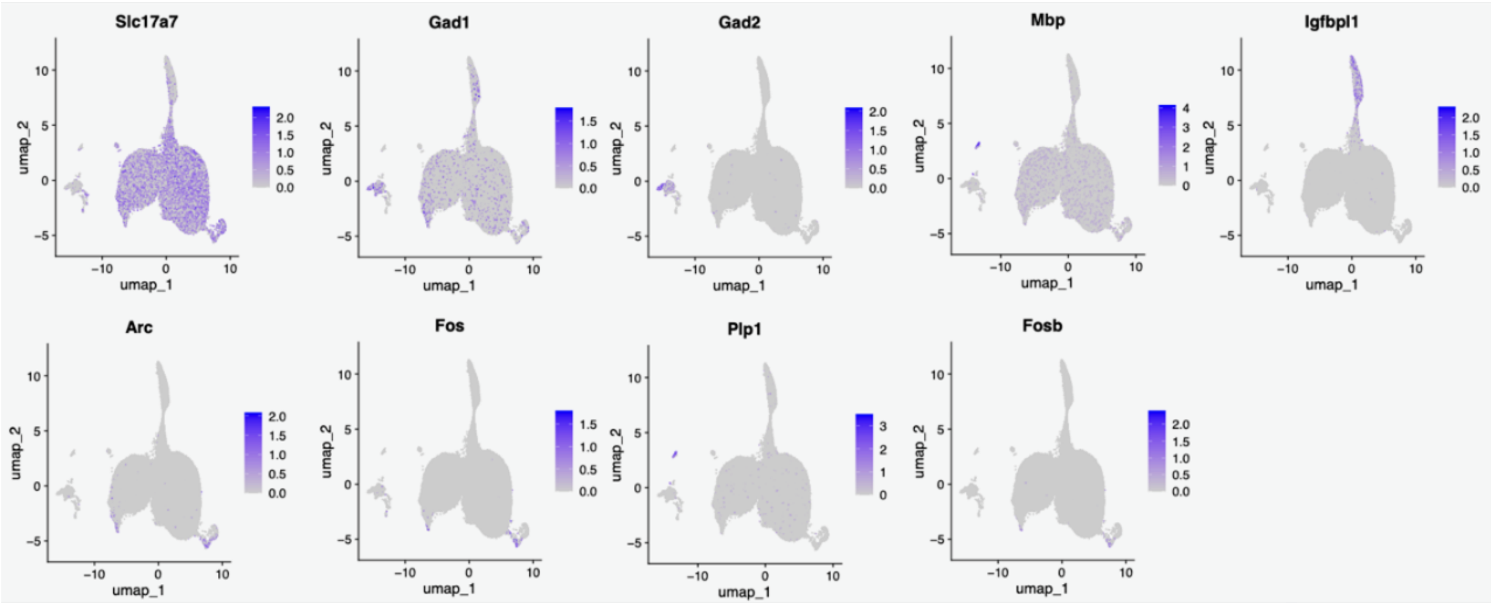


**Supplemental Figure 7**. Cluster markers for dentate granule cells. **A.** Marker gene expression in each cluster. Granule cells, Slc17a7; Interneurons, Gad1/2; Oligodendrocytes, Mbp, Plp1; Neuroblast, Igfbpl1; Activated granule cells, Arc, Fos, FosB. **B**. Distribution of individual marker genes in all Ext nuclei UMAP.

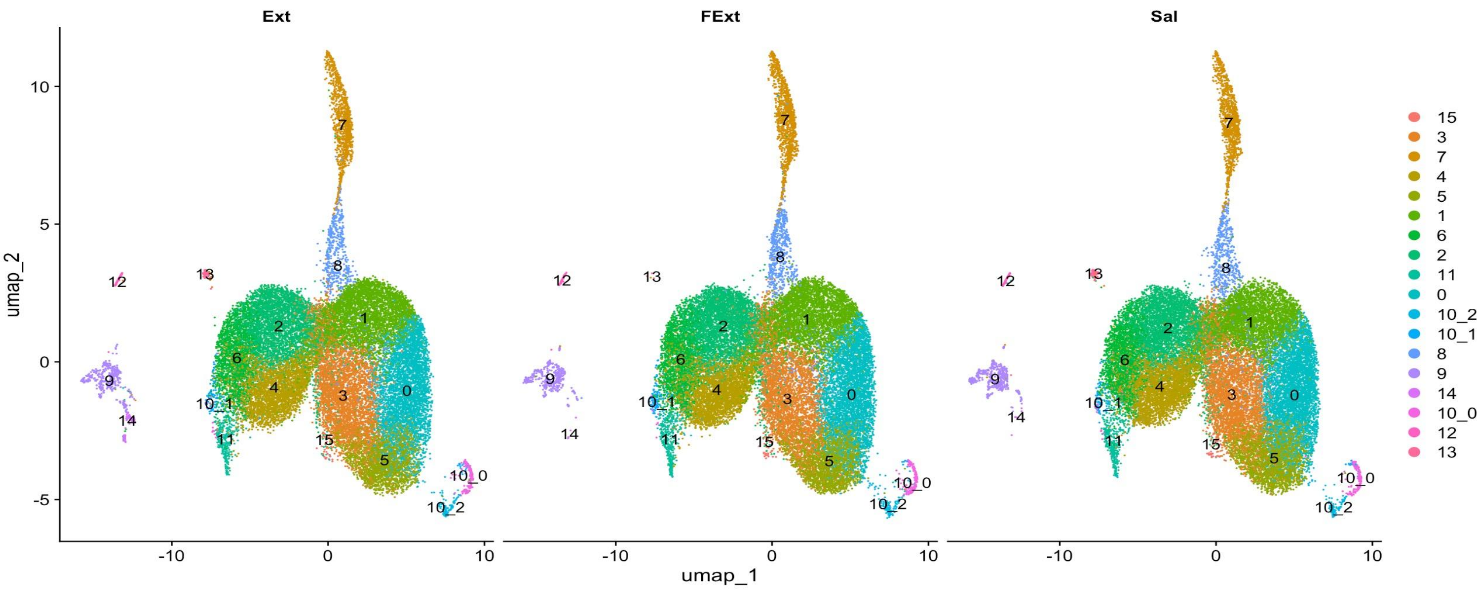


**Supplemental Figure 8**. UMAPs of S-Ext, F-Ext, and Ext-saline dentate granule cells.

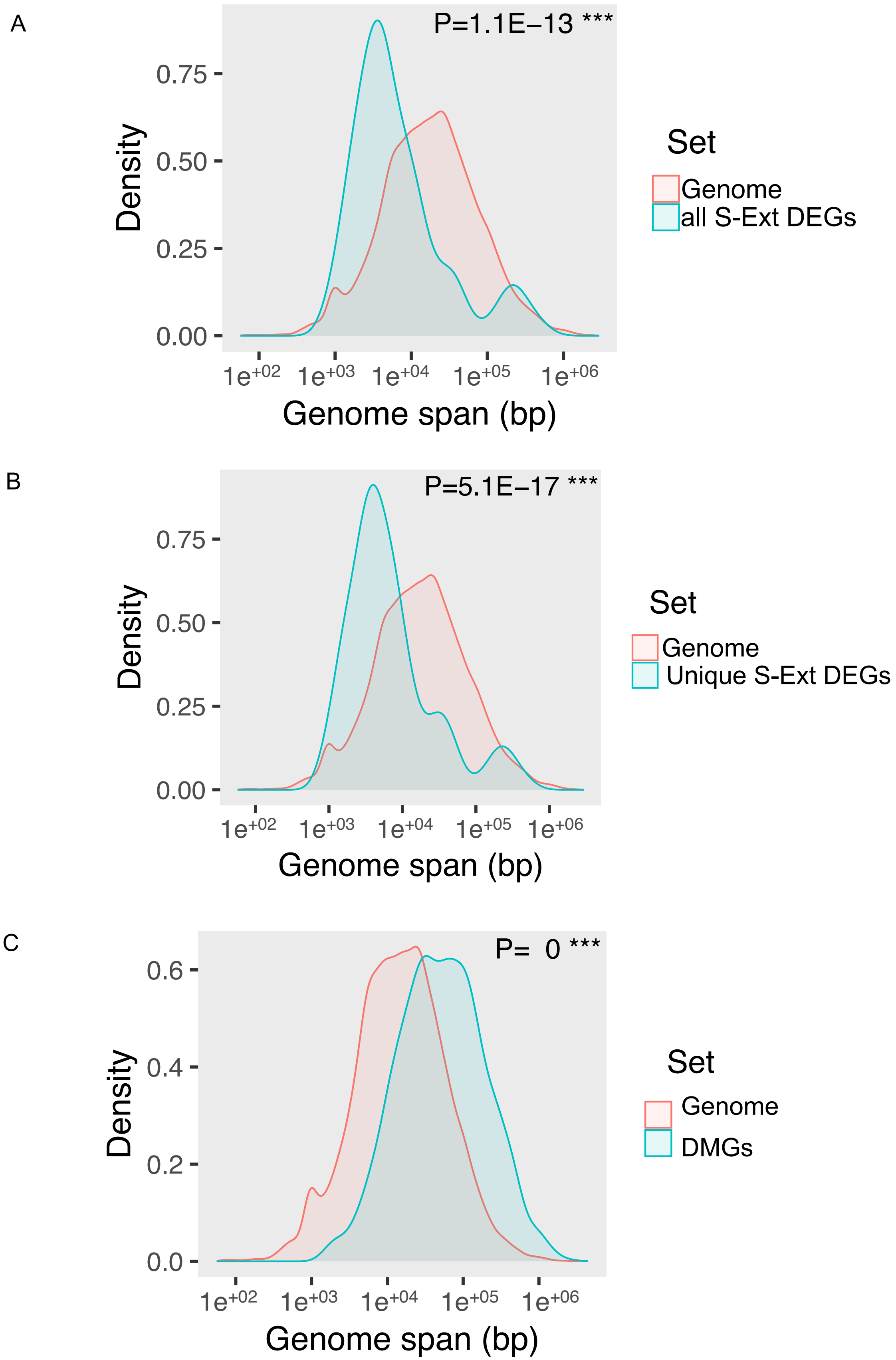


**Supplementary Figure 9**. Genome size of all and unique S-Ext DEGs in comparison to the genome average (A, B) and to the size of all Ext-DMGs (C).
